# *Drosophila* Toll links systemic immunity to long-term intestinal function

**DOI:** 10.1101/248138

**Authors:** Magda L. Atilano, Marcus Glittenberg, Shivohum Bahuguna, Lihui Wang, Petros Ligoxygakis

## Abstract

The intestine is an organ where immune and metabolic functions are co-ordinated with tissue renewal via progenitor somatic stem cells (PSSCs). How this is achieved is still unclear. We report that in *Drosophila*, a generalised infection increased PSSC numbers. This was mimicked by expressing a constitutive form of the immune receptor Toll in PSSCs and blocked when Toll was silenced via RNAi. Without infection, absence of bacterial recognition and downstream Toll signalling resulted in a short lifespan and an age-dependent decrease of PSSCs and gut microbiota. The latter implied a metabolic environment incompatible with the presence of bacteria. Indeed, infection or constitutive Toll signalling in PSSCs triggered 4E-BP transcription in enterocytes, while loss of signalling reduced it. 4E-BP controlled fat levels and sustained the microbiota suggesting that Toll-dependent regulation of 4E-BP was important for long-term gut function. Therefore, the Toll pathway is crucial for responses to both infection and microbiota.

## Introduction

Innate immunity is the first-line host defence conserved in all metazoans and plants (reviewed in Ronald and Beutler, 2010). In this context, Toll-like receptor (TLR) signalling is one of the most important mechanisms by which the innate immune system senses the invasion of pathogenic microorganisms in both mammals and Drosophila. Unlike its mammalian counterparts however, the fruit fly Toll is activated by an endogenous cytokine-like ligand, the Nerve Growth Factor homologue, Spz (Weber et al, 2003). Spz is processed to its active form by the Spz-Processing Enzyme (SPE) (Jiang et al, 2006). Two serine protease cascades converge on SPE: one triggered by bacterial or fungal serine proteases and a second activated by host receptors that recognise bacterial or fungal cell wall. Prominent among these host receptors is the Peptidoglycan Recognition Protein-SA or PGRP-SA (Michel et al, 2001), which is considered to preferentially bind to Lys-type Peptidoglycan from Gram-positive bacteria (Chang et al, 2004). When the recognition signal reaches the cell surface, it is communicated downstream via the Toll receptor and a membrane-bound receptor-adaptor complex including dMyd88, Tube (as an IRAK4 functional equivalent) and the Pelle kinase (as an IRAK1 functional homologue) (Marek and Kagan 2012). Transduction of the signal culminates in the phosphorylation of the IκB homologue, Cactus (Daigneault et al, 2013). This modification requires the fly βTrCP protein Slimb (Daigneault et al, 2013) and targets Cactus for degradation, leaving the NF-κB homologue DIF to move to the nucleus and regulate hundreds of target genes including a battery of powerful antimicrobial peptides (AMPs) (de Gregorio et al, 2002).

Similar to mammals, the high capacity of intestinal epithelial regeneration in *Drosophila* depends on intestinal stem cells (ISCs). These multipotent ISCs, which are distributed along the basement membrane of the posterior midgut have a simple pattern of division (Ohlstein and Spradling 2006; Ohlstein and Spradling 2007, reviewed in Lemaitre and Miguel-Aliaga, 2013). An ISC divides to produce itself and an enteroblast (EB), which will undergo terminal differentiation into an enterocyte (EC) or an enteroendocrine cell (EE). Progenitor cells (ISCs and EBs) express a transcription factor called Escargot (esg) (Ohlstein and Spradling 2006; Ohlstein and Spradling 2007; Amcheslavsky et al 2009). Thus, expression of *esg* is often used as a surrogate marker for studying both ISCs and EBs in the anterior midgut. Previous studies have shown that direct local damage to the gut by oxidative stress, toxins or ingestion of bacteria leads to EC apoptosis, stimulating ISCs to proliferate and replenish EC numbers (Amcheslavsky et al 2009; Buchon et al, 2009). In this context, intestinal microbiota impacted on the rate of ISC proliferation by stimulating them to keep a higher “baseline” of epithelial turnover (Buchon et al, 2009). However, less is known about what happens after systemic infection. Septic injury with the Gram-negative bacterium *Erwinia carotovora carotovora-15 (Ecc-15)* promotes ISC proliferation through cytokine signalling mediated by blood cells (Chakrabarti *et al*, 2016). In addition, sterile wounding also causes EC apoptosis and triggers ISC proliferation (Takeishi et al, 2013).

The intestine has also a metabolic role and this depending on the context is coordinated by insulin, FOXO and TOR (reviewed in Miguel-Aliaga *et al,* 2018). One of the downstream components of this signalling network is the translational inhibitor 4E-BP (Miron et al, 2003). It has been hypothesised that the control of 4E-BP activity in flies could provide a means to control fat metabolism during stress conditions (Teleman et al, 2005). In adults, 4E-BP is important for infections that activate the Toll pathway, since 4E-BP mutant flies are highly susceptible to fungal and Gram-positive bacterial infections (Bernal and Kimbrell 2000). In larvae, FOXO induces 4E-BP activation following *Ecc-15* oral infection or starvation stress, which tips the balance towards translation of cap independent transcripts including AMPs (Vasuvedan *et al*, 2017). Thus, 4E-BP may provide the link between systemic Toll activation and metabolic control.

After the discovery of intestinal T-cells expressing TLRs, we know that the immune system can discriminate between and regulate appropriate responses towards, non-infectious non-self (e.g. gut microbiota) vs. infectious non-self (reviewed in Kubinak and Round 2012). However, it is less clear how immune response signals from a systemic infection will be integrated at the intestine level by ISCs and ECs with the aim of maintaining tissue integrity and normal gut microbiota. In the present work, we investigated how systemic infection triggering Drosophila Toll/TLR signalling may be linked to long-term intestinal homeostasis.

## Results and discussion

To trace intestinal progenitor cells (ISCs and EBs) we used the GAL4/GAL80^ts^/UAS system (Suster et al, 2004). The driver for GAL4 expression was the *esg* gene promoter. Normally, GAL80 is an inhibitor of GAL4 but the temperature sensitive allele used was able to do this only at the permissive temperature (18°C) and not at the restrictive temperature (30°C). In all the experiments described in this work, developing flies were cultured at 18°C and on eclosion emerging adults were shifted to 30°C. When 20-day old adults expressing GFP under the *esg* promoter in the anterior midgut (Fig. 1A) were injected in the thorax with the opportunistic fungal pathogen *Candida albicans* (*C. albicans*), we observed a statistically significant increase of GFP-positive cells 36h post-infection (Fig. 1B). This result indicated that systemic immunity regulated intestinal progenitor cell numbers. Since Toll is the pathway primarily responding to fungal infections in Drosophila (reviewed in Kounatidis and Ligoxygakis 2012), we attempted to reproduce the result by mimicking Toll triggering by infection. To this end, we expressed Toll10B, a constitutively active form of Toll (Shia *et al,* 2009) in *esg*-expressing cells with the GAL4/GAL80^ts^/UAS system (*esg*^*ts*^>*Toll10B*). This led to a significant increase of GFP positive (GFP^+^) cells (Fig. 1C). When we expressed UAS-Toll10B and UAS-GFP with *Su(H)-*GAL4, GAL80^ts^ [*Su(H)^ts^*, an EB-specific GAL4; Zeng et al, 2010], we also observed a significant rise in the numbers of large GFP^+^ cells (resembling EBs) in *Su(H)^ts^>Toll10B* flies (Fig. 1C). This could be interpreted as either 1) Toll signalling creating a “roadblock” for further EB differentiation (hence the increase in GFP cells may be a “backlog” of EBs) or 2) Toll inducing division of the existing EBs to produce more of the same. However, staining with an antibody against the phosphorylated form of Histone 3 (pH3) to assay cell proliferation did not show any pH3 staining in *Su(H)^ts^>Toll10B* guts (Fig. S1A). This indicated that the increase of GFP^+^ cells in *Su(H)^ts^>Toll10B* flies was not a result of EB proliferation. Nevertheless, data in Fig. 1 show that systemic *C. albicans* infection or constitutive Toll signalling was sufficient to increase the pool of progenitor cells.

**Figure 1.**
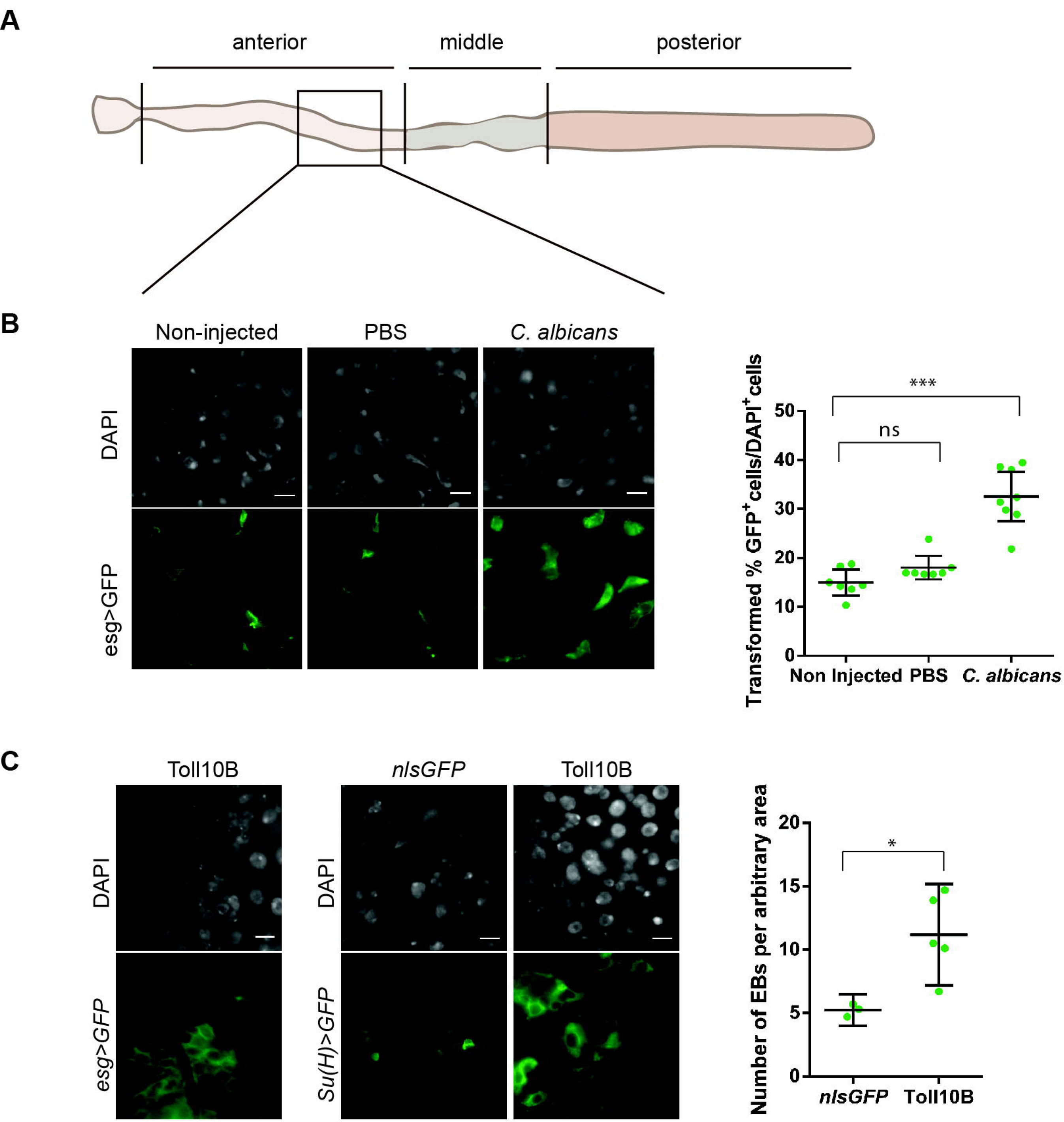
Intestinal progenitor cells respond to systemic C. albicans infection. Focusing on an area of the anterior midgut close to the border with middle midgut **(A)** we found a significant increase in large GFP-positive cells (green) found in *esg*^*ts*^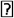GFP intestines following injection by *C. albicans* in the thorax compared to sterile injury with PBS **(B)**. This was mimicked in the absence of inf ection or injury by the expression of a constitutively active form of the Toll receptor (Toll10B) when expressed in progenitor cells (*esg-Gal4*) or just in EBs [*Su(H)-Gal4*] and compared to expression of nlsGFP **(C)**. This implies that the large GFP cells seen in **(B)** were EBs. All nucleic were stained with DAPI (grey). In the plots, each cicle represents the counted area from a single gut; 95% confidence interval is displayed (* (p<0.05, *** p<0.001, ns = non-significant).

We next asked whether, in addition to being sufficient, Toll was also necessary for controlling intestinal epithelial renewal following systemic fungal infection. We silenced Toll (using an RNAi line from the VDRC collection) in progenitor cells using *esg*^*ts*^. Development proceeded at the permissive temperature (18°C, GAL80 ON, GAL4 OFF, Toll RNAi OFF). Emerging day 1 adults were transferred to the restrictive temperature of 30°C (GAL80 OFF, GAL4 ON, Toll RNAi ON) and infected with *C. albicans* 20 days later. After infection we examined the number of GFP-positive cells at 36h post-infection. As a control for the RNAi mechanism we used a random line from the same VDRC collection (*UAS-CG7923^RNAi^*) that did not compromise host survival when infected with *C. albicans* and exhibited a normal lifespan compared to its genetic background in the absence of infection (our unpublished observations). The use of an endogenous gene as a control instead of an inert one (e.g. RFP) ensured that we controlled for the actual triggering of the RNAi mechanism *in an existing host gene*.

In control flies, we observed an increase of GFP-positive cells when comparing sterile injury (PBS) with septic injury (*C. albicans*) (Fig. 2A), which was statistically significant when quantified (Fig. 2B). Silencing Toll prevented an increase in progenitor cells following infection since GFP-positive cells after PBS injection or *C. albicans* challenge were statistically inseparable when compared (Fig. 2B). In contrast to controls, *esg*^*ts*^>*Toll*^*RNAi*^ flies were unable to increase ISC proliferation following infection (Fig. S1B). Taken together these results indicated that Toll was also necessary for the long-term renewal of the intestinal epithelium following infection. Moreover, we noted that the number of GFP positive cells in *esg*^*ts*^>*CG7923*^*RNAi*^ flies was significantly higher than in *esg*^*ts*^>*Toll*^*RNAi*^ flies after PBS treatment, raising the possibility that the intestines of *esg*^*ts*^>*Toll*^*RNAi*^ had less progenitor cells in 20-day old adult guts even in the absence of infection (Fig. 2B).

**Figure 2.**
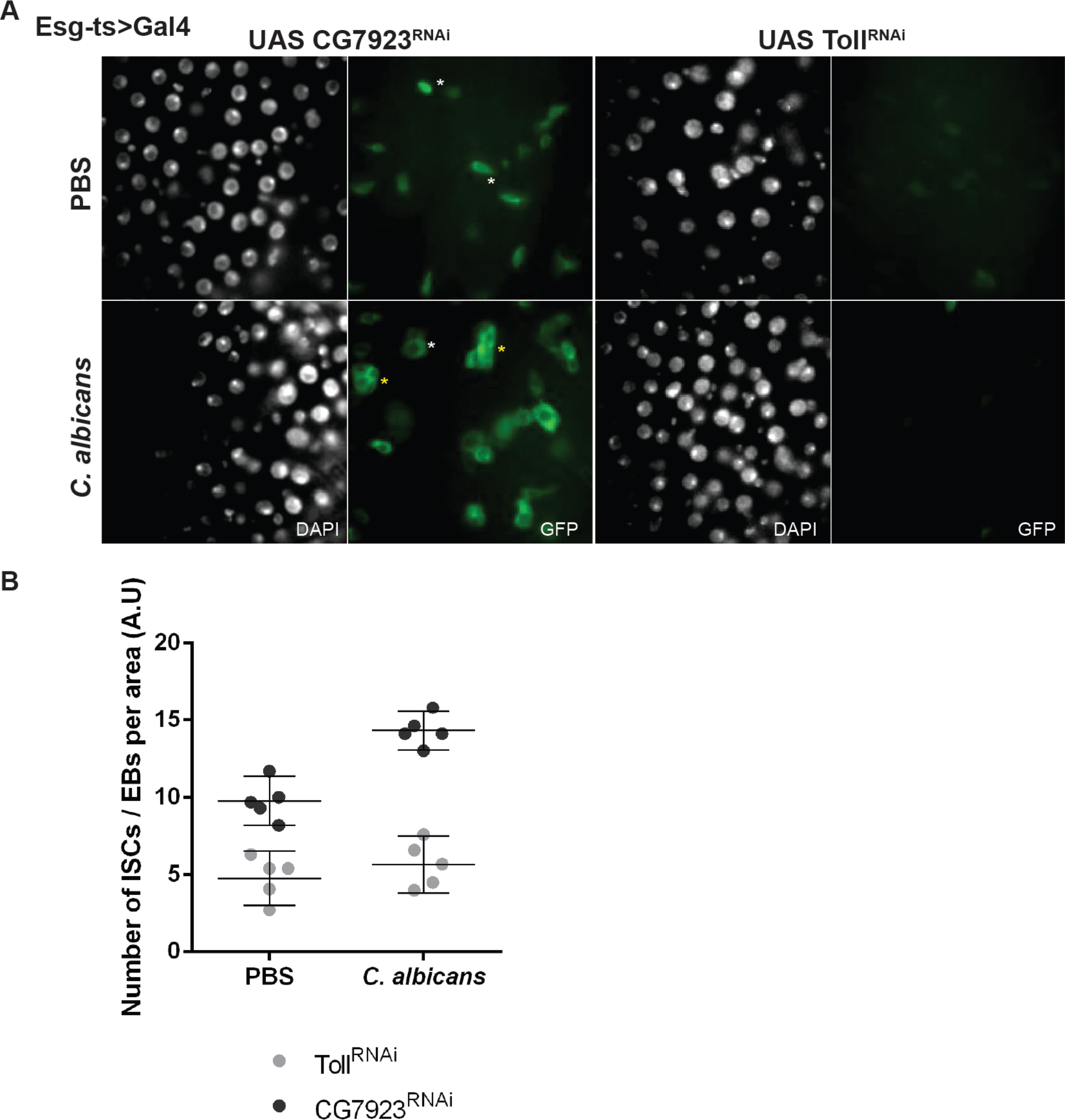
Silencing Toll in intestinal progenitor cells prevents their increase during systemic infection. **(A)** 36 hours following PBS injection or systemic *C. albicans* infection the number of ISCs / EBs (marked with GFP, green) increases (UAS-CG7923 RNAi), but not when the function of Toll is reduced (UAS-Toll^RNAi^) specifically in these cells (*esg*^*ts*^>Gal4) **(B)**. Systemic infection may also induce enhanced clustering (yellow asterisks) and morphological changes (compare white asterisks) to the ISCs / EBs, which are diminished when Toll receptor activity is reduced. All nuclei stained with DAPI (grey). Each circle represents a counted area from a single gut; 95% confidence interval is displayed.

To test long-term renewal of the intestinal epithelium in the absence of infection, we assayed guts from 6-day old and 20-day old *esg*^*ts*^>*Toll*^*RNAi*^ flies and compared them to chronologically age-matched *esg*^*ts*^>*CG7923*^*RNAi*^ controls. When comparing 6-day old and 20-day old flies, we found that at 20 days the shape of GFP positive progenitor cells in guts of *esg*^*ts*^>*Toll*^*RNAi*^ adults was significantly altered compared to *esg*^*ts*^>*CG7923*^*RNAi*^, with cells becoming smaller and more rounded (Fig. 3A). They were also significantly reduced in numbers (Fig. 3B), with the tissue becoming extremely fragile to handle. Moreover, cultivable intestinal microbiota was significantly lower at 20-days as measured by Colony Forming Units (CFUs) (Fig. 3C). The same effect on cell numbers and cell shape in 20-day old flies was also observed when Toll was silenced in EBs [*Su(H)ts>Toll^RNAi^*] (Fig. S2).

**Figure 3.**
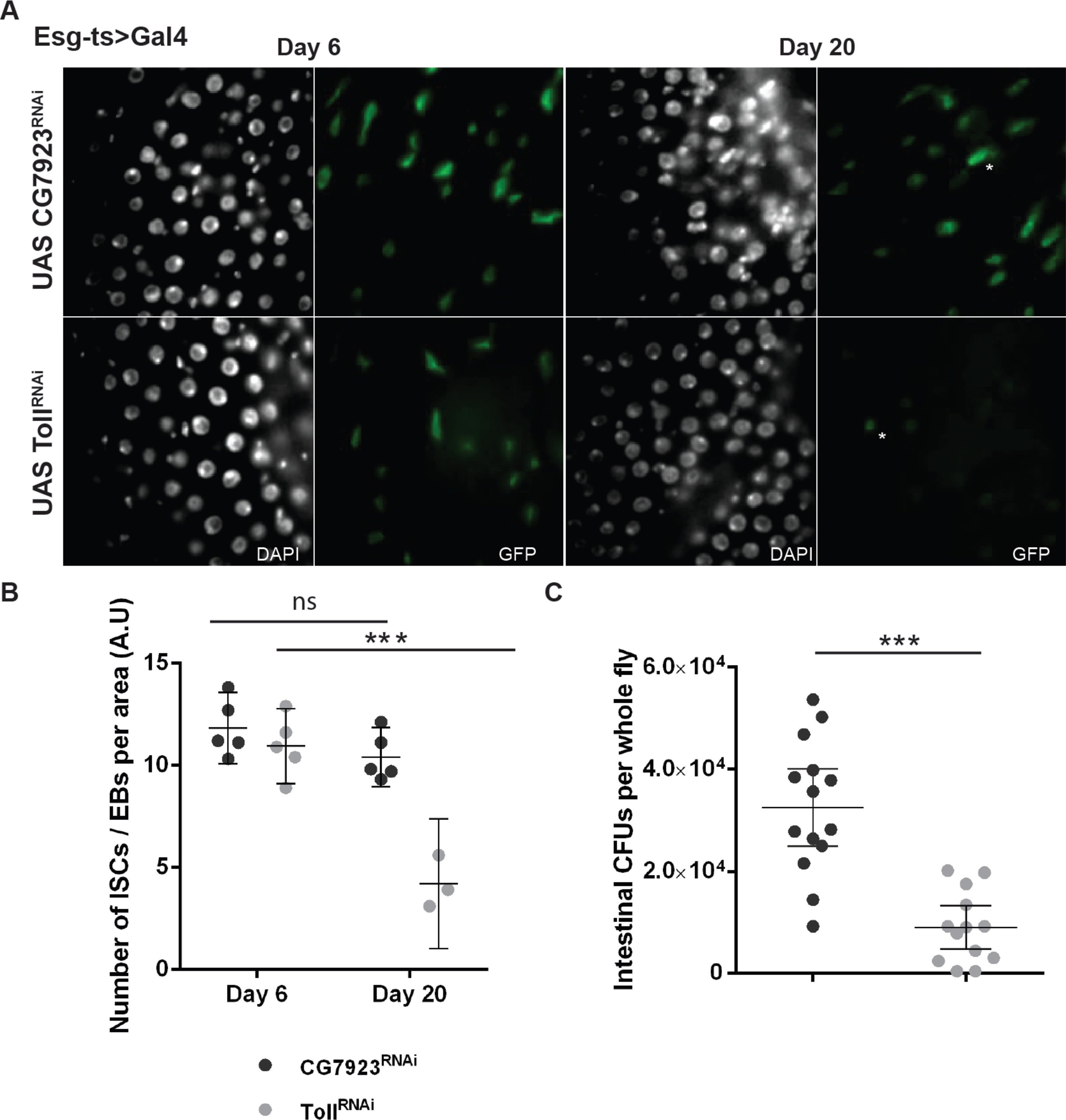
The Toll receptor influences the long-term renewal of gut progenitor cells in the absence of infection. **(A)** RNAi knockdown of the UAS-CG7923 was randomly chosen from the VDRC collection as a control for the RNAI effect. Using UAS-Toll^RNAi^ (from the same VDRC collection) the Toll receptor was knocked-down specifically in ISCs and EBs (esg-ts > Gal4) of the adult gut. This led to altered morphology of progenitors, where the cells generally remained round, rarely adopting the characteristic elongated and irregular shape often observed with ISCs/EBs a reduction in cell number and alters their morphology, where the cells generally remain rounded, rarely adopting the characteristic elongated and irregular shape often observed with ISCs / EBs (compare cells marked with white asterisks). **(B)** There was also a significant reduction in the numbers of progenitor cells when Toll is knocked-down [day 6 compared with day 20 - *** (p<0.001)]. Each dot in the graph represents a counted area from a single gut taken from 3 biological repeats **(C)** Toll knocked-down resulted in a significant reduction of cultivable intestinal microbiota (CFUs) (p<0.001) in 20-day old flies; 95% confidence intervals are displayed.

In the absence of infection, 20-day old flies mutant for the upstream-most Toll pathway component namely, *PGRP-SA^seml^* (Michel *et al*, 2001) showed a reduction in ISC proliferation with fewer ISCs dividing to EBs (Fig 4A). The latter were marked with an antibody against anti-hrp (Han *et al,* 2015; O’Brien *et al,* 2011). Moreover, PGRP-SA-deficient flies exhibited a significant reduction in the numbers of progenitor cells in general, and ISCs in particular (Fig. 4B). This was also reflected in the very low ISC proliferation seen i n *PGRP-SA^seml^* mutants compared to controls (Fig. S3). In addition, *spz* mutant flies had significantly reduced ECs (Fig. S4). Expression of Toll10B in *PGRP–SA^seml^* progenitor cells was able to reconstitute progenitor cells in the anterior midgut (Fig. S5A) with numbers statistically indistinguishable from yw, the genetic background of *PGRP-SA^seml^* (Fig. S5B). Moreover, 20-day old PGRP-SA and DIF mutants showed a reduction in intestinal microbiota when total intestinal 16S rRNA genes were semi-quantified (Fig. S6), as well as significantly reduced lifespan of both male and female flies (Fig. S7). Taken together, these results showed that Toll was both necessary and sufficient for regulating the long-term regeneration potential of the intestine after an immune challenge but also in the absence of infection.

**Figure 4.**
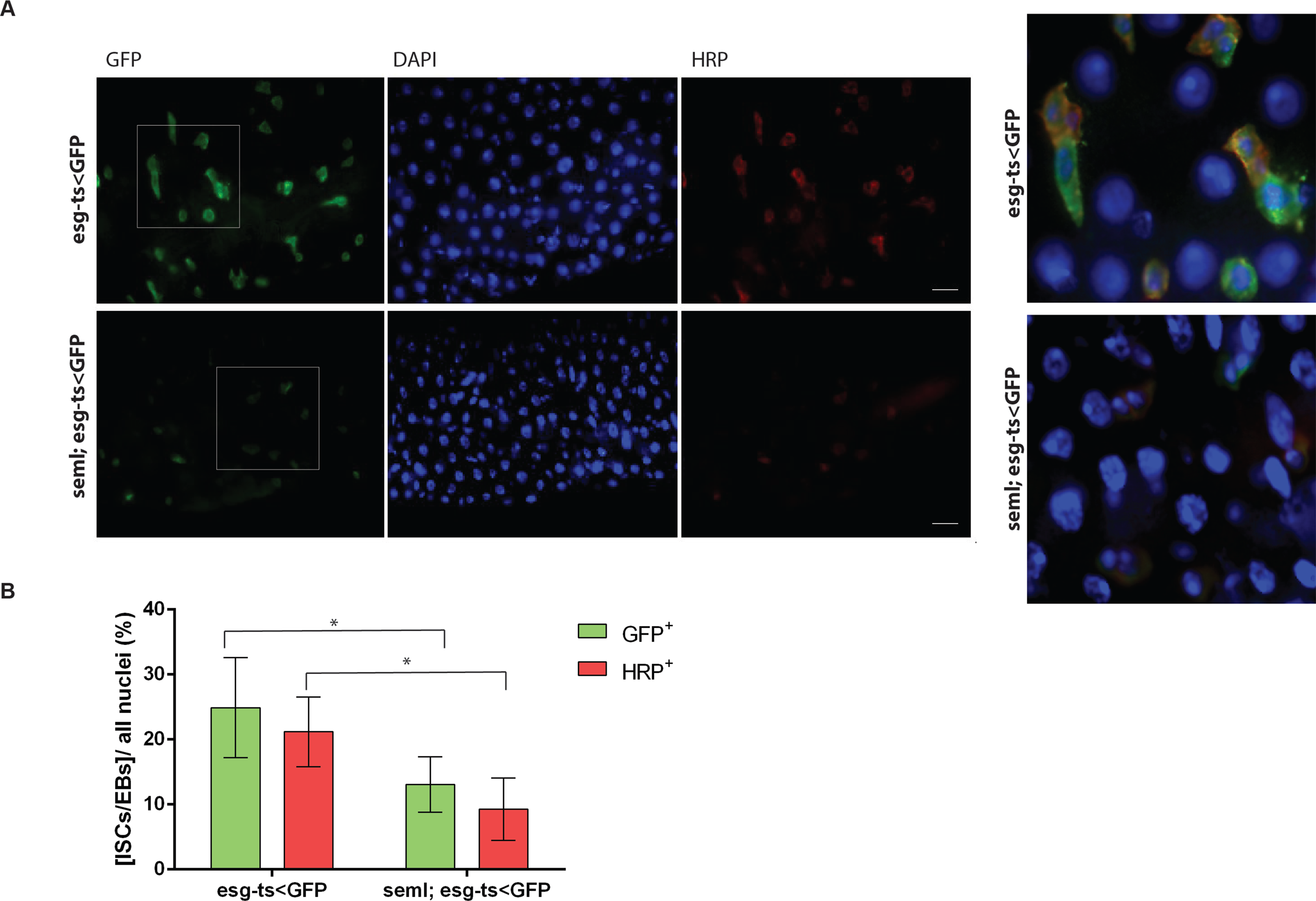
PGRP-SA^*seml*^ mutant flies have less intestinal progenitor cells. **(A)** In the absence of infection, ISC (HRP positive, GFP positive) divide to produce EBs (HRP negative, GFP positive). However, in 20-day old flies that were deficient for PGRP-SA this division was not observed (see also insets). **(B)** Quantification of progenitor cells and ISCs showed that these were significantly reduced (*p<0.05; error bars display 95% confidence intervals, guts from 4 biological repeats were analysed). GFP expression was directed by the UAS dependent mCD8GFP transgene, which marked the cell membranes of the progenitor cells including ISCs and EBs (DAPI all nuclei).

Microbiota reduction pointed towards a metabolic shift in the gut that was unable to sustain the normal density of bacterial populations in the absence of PGRP-SA/Toll signalling. The *Drosophila* homologue of 4E-BP has been shown to be activated by infections that trigger Toll signalling and has NF-κB binding sites in its promoter (Bernal and Kimbrell, 2000). Such infections include *C. albicans* and 4E-BP null mutants are susceptible to this particular immune challenge (Levitin et al, 2007). Induction of 4E-BP activity can shift the balance towards translation of uncapped mRNAs including those of AMPs (Kang *et al,* 2017) and decelerate fat metabolism (Teleman *et al,* 2005). Indeed, *C. albicans* infection of 20-day old flies induced a 4E-BP transcription reporter construct (Bernal and Kimbrell 2000) in ECs but not in progenitor cells (Fig 5A). This induction was significantly higher than injection of sterile PBS or the non-infected control (homeostasis) (Fig. 5B). 4E-BP transcriptional induction was also observed following *Staphylococcus aureus* (*S. aureus*) infection again in ECs (Fig. 6A). After immune challenge with *S. aureus*, the number of GFP+ marked progenitors were significantly increased as was transcription of 4E-BP (Fig. 6B). Importantly, Toll10B expression in progenitor cells emulated the above as it resulted also in the transcriptional upregulation of 4E-BP in ECs in a manner comparable to *C. albicans* as well as *S. aureus* infection (Fig. 7A). This increase was significantly higher than the non-infected control (Fig. 7B).

**Figure 5.**
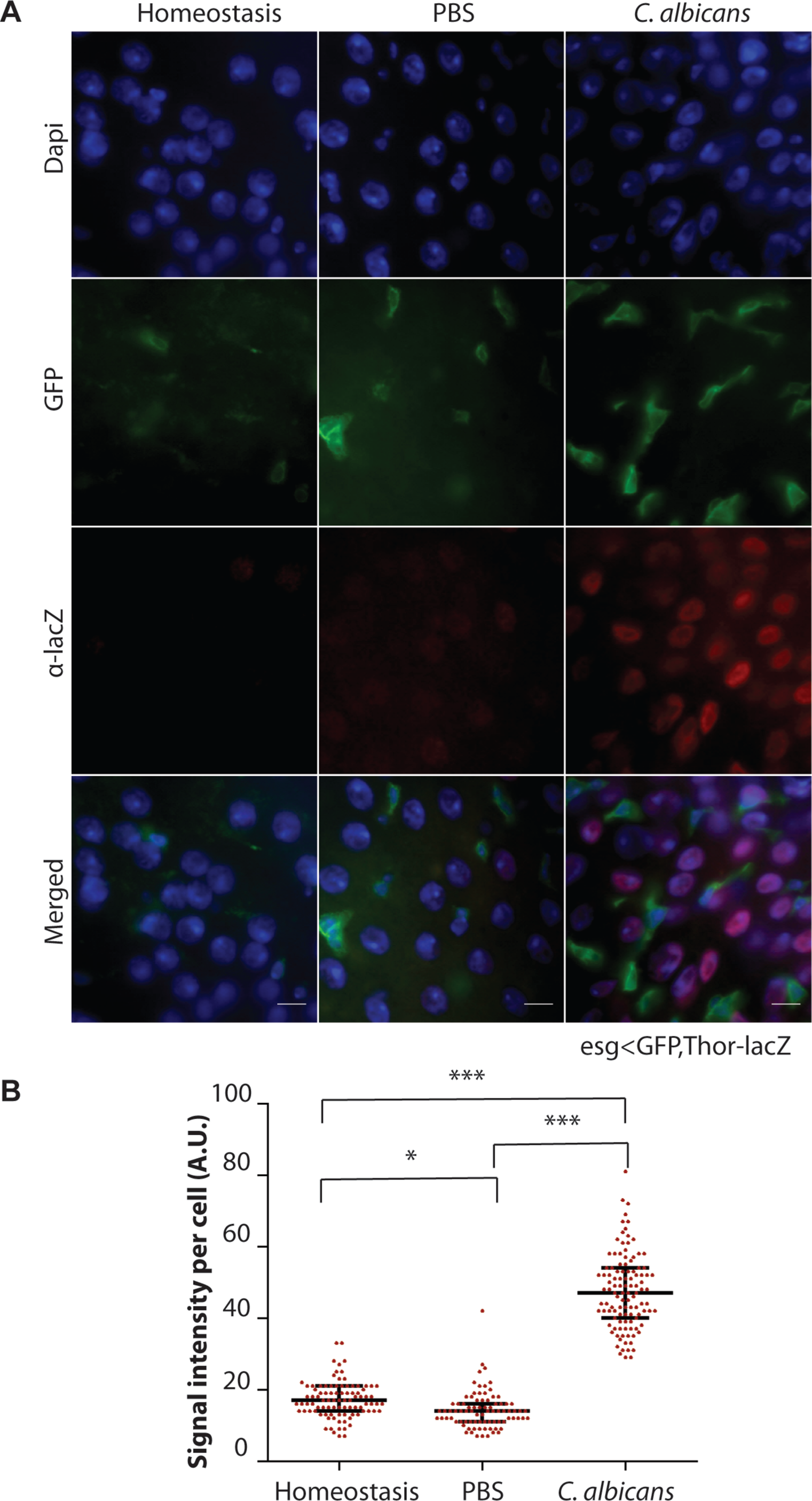
Systemic *C. albicans* infection activates d4E-BP (Thor) transcription in enterocytes. (A) *thor-lacZ; esg^ts^>GFP* flies injected with *C. albicans* were sampled 36 hours post-infection and compared to non-treated (homeostasis) or those injected with PBS (sterile injury). Gut cells stained with DAPI (blue), anti-β-galactosidase (red) and anti-GFP expressing cells (marking both ISCs and EBs). Shown are representative images from the anterior midgut taken at 63x. (B). Quantification of *thor-lacZ* expression upon systemic infection. Intensity measured using ImageJ, subtraction of the background was performed for all the samples. Ten guts were analysed (approximately 50 cells analysed per gut), 95% confidence intervals displayed, *p<0.05, *** p<0.001.

**Figure 6.**
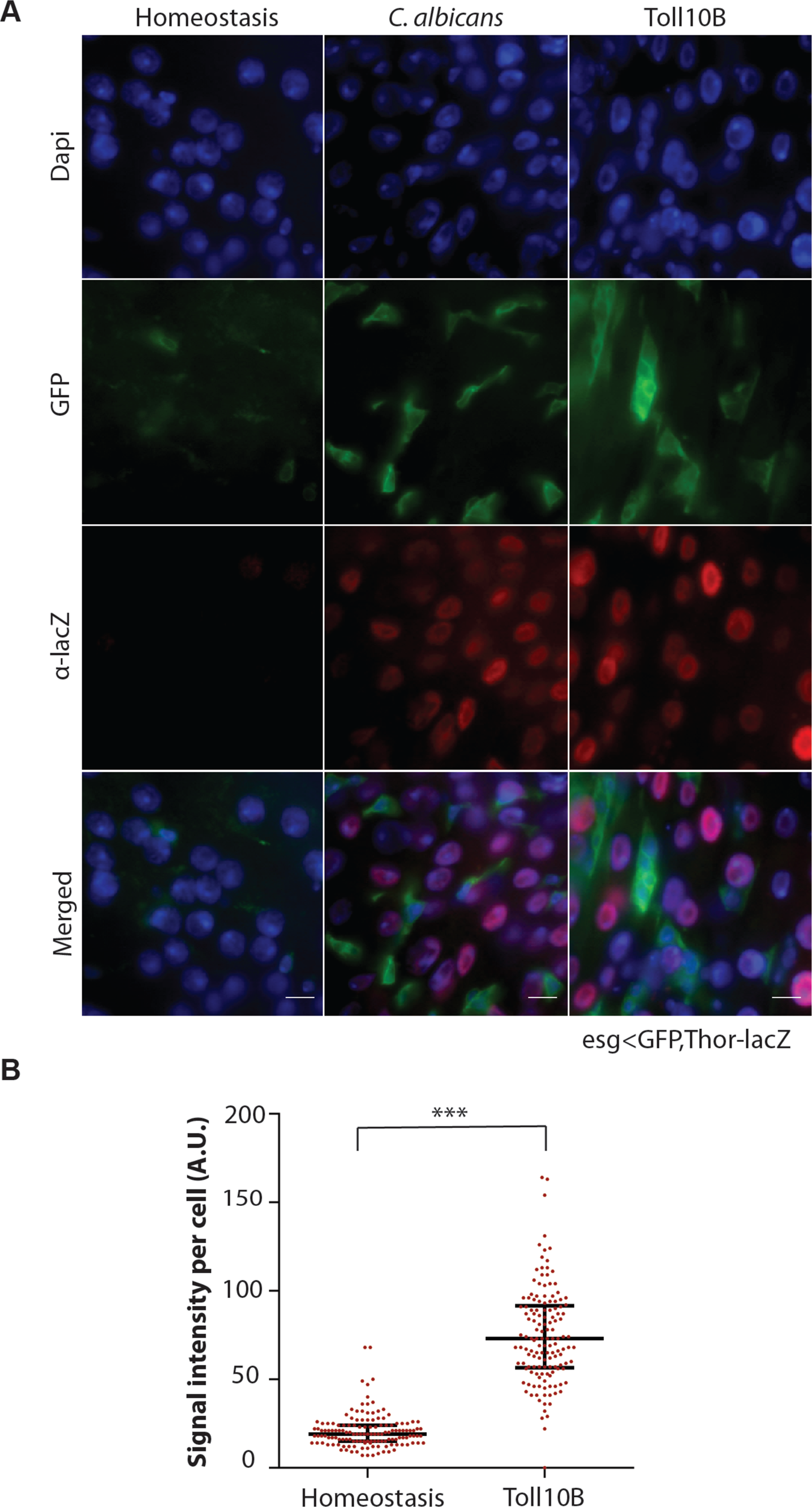
Systemic infection of S. aureus activates d4E-BP transcription in enterocytes. **(A)** *thorlacZ; esg^ts^>GFP* flies injected with S. aureus were sampled 36 hours post-infection and compared to non-treated or those injected with PBS (sterile injury). Gut cells stained with DAPI (blue), anti-β-galactosidase (red) and anti-GFP expressing cells (marking both ISCs and EBs). Shown are representative images from the anterior midgut taken at 63x. **(B)** Quantification of progenitor cells following *S. aureus* infection indicating that systemic bacterial infection can also increase the population of intestinal progenitor cells **(C)** Quantification of *thor-lacZ* expression upon systemic *S. aureus* infection. Intensity measured using ImageJ and subtraction of the background was performed for all the samples. Ten guts were analysed (approximately 50 cells analysed per gut), 95% confidence intervals displayed, *p<0.05, *** p<0.001.

**Figure 7.**
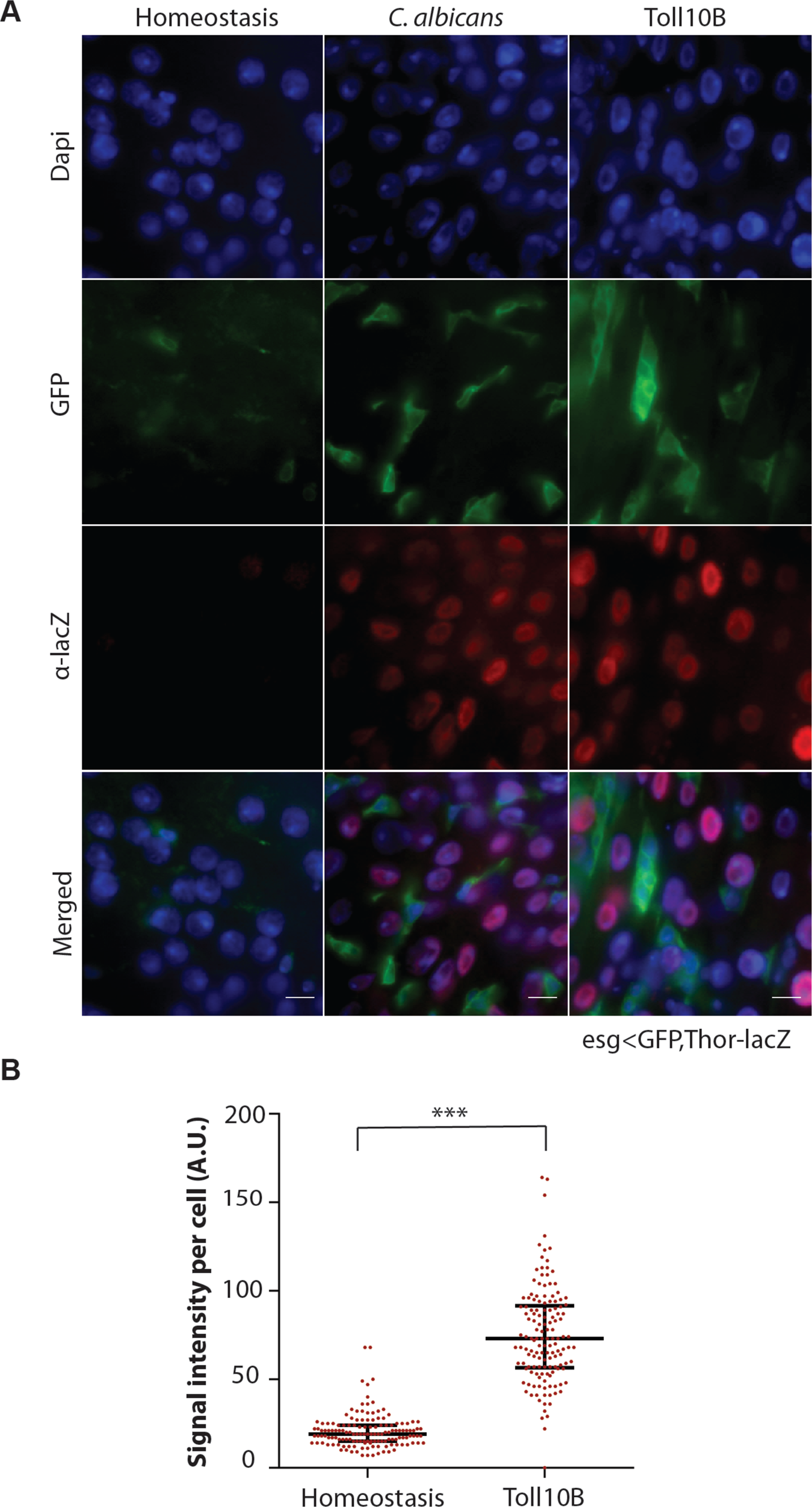
Toll10B expressed in progenitor cells activates d4E-BP transcription in enterocytes. **(A)** Flies injected with *C. albicans* were sampled 36 hours post-infection and compared to non-treated (homeostasis) or those expressing Toll10B. Gut cells stained with DAPI (blue), anti-β-galactosidase (red) and anti-GFP expressing cells (marking both ISCs and EBs). Shown are representative images from the anterior midgut taken at 63x. Two left hand columns show *thor-lacZ* **(B)**. Quantification of *thor-lacZ; esg^ts^>GFP*; right column *thor-lacZ; esg^ts^>GFP* flies also express Toll10B transgene. **(B)** Quantification of thor-lacZ expression upon systemic infection. Intensity measured using ImageJ, subtraction of the background was performed for all the samples. Ten guts were analysed (approximately 50 cells analysed per gut), 95% confidence intervals displayed, *** p<0.001.

Increase of 4E-BP transcription had an effect on fat metabolism as reported previously (Teleman *et al*, 2005). 48h after *C. albicans* infection, 20-day old control flies had elevated fat levels when normalised to total body protein (Fig. 8). This increase became more prominent when considering intestinal-only triglycerides (TGs) (Fig. S8). This indicated that systemic infection in *Drosophila* adults resulted in increased intestinal accumulation of TGs and increased systemic levels of fat. Infection of 4E-BP null mutants or flies with silenced 4E-BP in ECs through the *NP1-GAL4, GAL80^ts^* (*NP1^ts^*>*4E-BP*) configuration did not produce this phenotype (Fig. 8, Fig. S8). Overexpression of Toll10B in progenitor cells had a similar effect as *C. albicans* infection but this was suppressed in a 4E-BP null genetic background indicating that fat levels were controlled by 4E-BP (Fig. 8, Fig. S8). In the absence of infection, loss of Toll signalling (*PGRP-SA^*seml*^* mutants) resulted in significantly reduced fat levels in 20-day but not in 5-day old flies (Fig. 9). The latter had the same levels of fat as the controls (Fig. 9). Reduction of fat was accompanied by a reduction in intestinal CFUs (Fig. S9). It has been established that TOR-mediated phosphorylation keeps 4E-BP activity suppressed (Hay and Sonenberg 2004). 20-day old PGRP-SA mutant flies with TOR-RNAi in ECs or treated with the mTOR inhibitor rapamycin restored fat quantities (Fig. 9) and CFUs (Fig. S9) to the level of the *yw* control.

**Figure 8.**
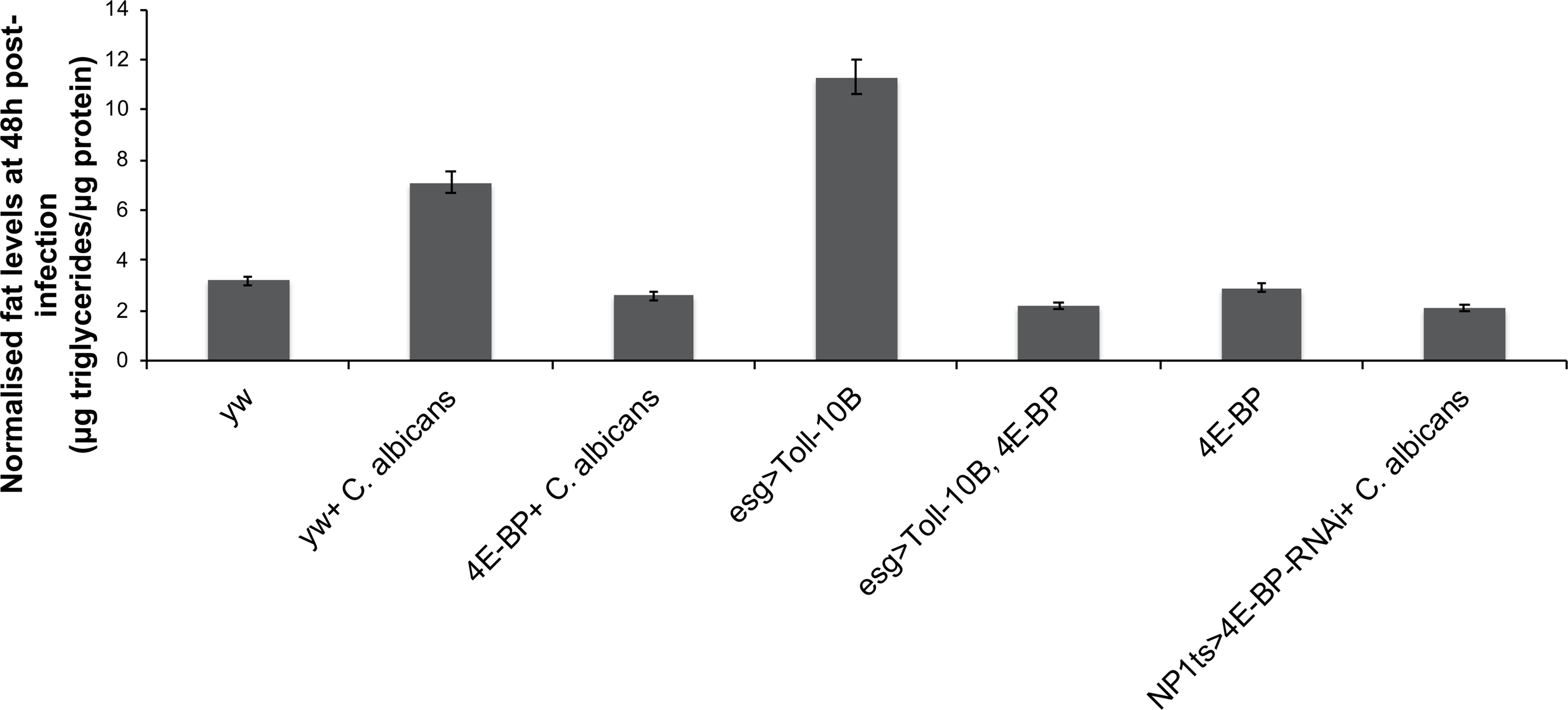
4E-BP regulates systemic fat levels following infection or Toll over-expression in intestinal progenitor cells. Following *C. albicans* infection systemic fat levels were increased (comparison between *yw* and yw-*C. albicans* p<0.001). The presence of d4E-BP ensured that fat levels are not “burnt” as fast during infection since Thor mutants (4E-BP-*C. albicans*) or flies with silenced 4E-BP in the ECs (*NP1^ts^>4E-BP-RNAi*) that were infected had significantly less fat (comparison between yw-*C. albicans* and 4E-BP or NP1ts>4E-BP-RNAi p<0.001). Toll overexpression in progenitor cells through *esg-GAL4* was able to increase fat levels (comparison between *yw-C. albicans* and Toll-10B is non-significant but between yw and Toll-10B p<0.001). However, in a genetic background mutant for 4EBP, Toll overexpression did not have an effect (comparison between *4E-BP-C. albicans* and Toll non-significant). Error bars represent standard deviation. Bars represent mean values from three independent experiments.

**Figure 9.**
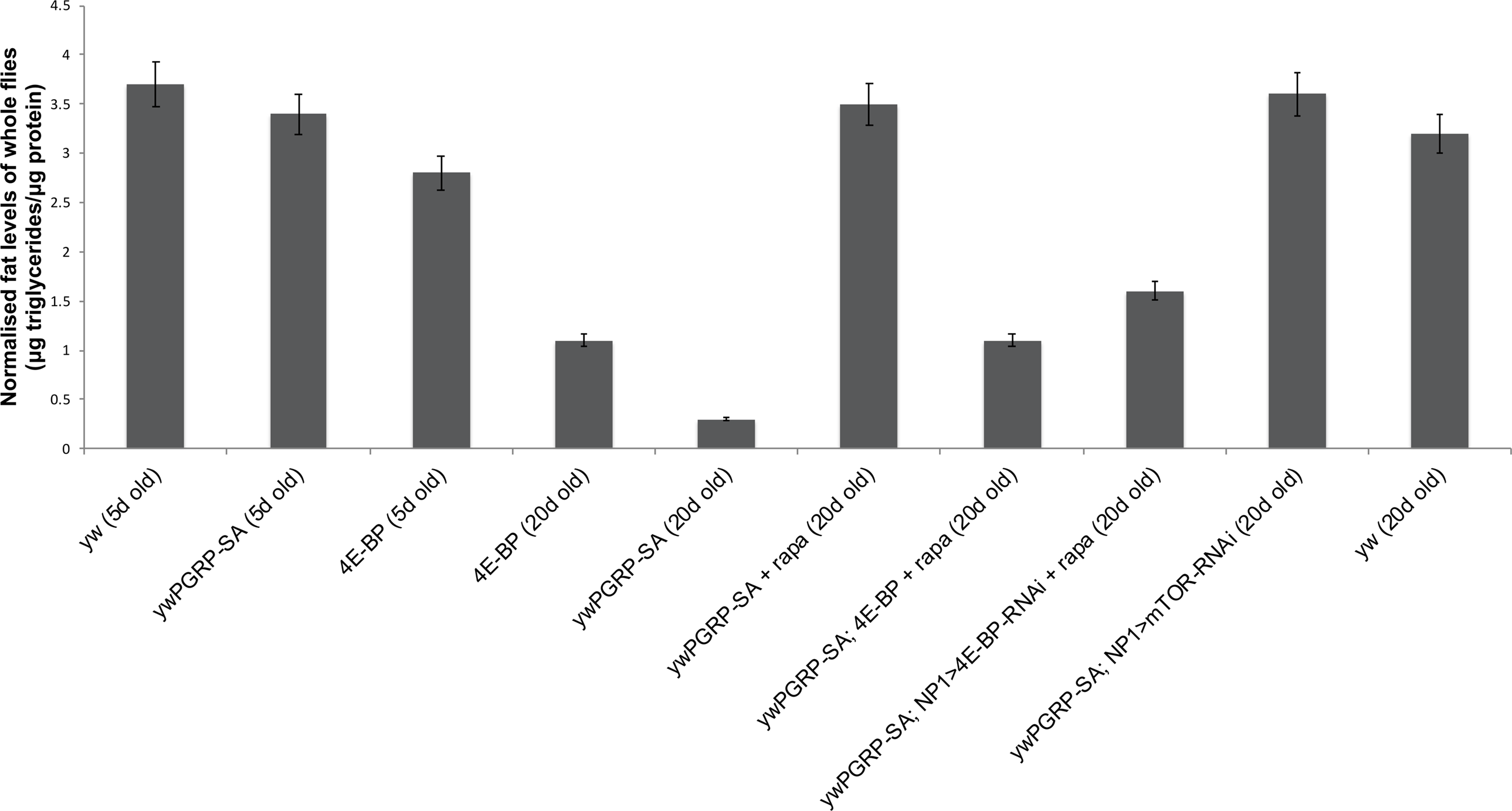
PGRP-SA regulates 4E-BP and fat content in an age-dependent manner. In the absence of infection, 20-day old (but not 5-day old) PGRP-SA mutants had significantly reduced fat levels (comparison between 20-day old *yw* and *PGPP-SA* p<0.001 but non-significant when comparing 5-day old flies). Lack of PGRP-SA increased mTOR activity since either treatment of *PGRP-SA* mutants with rapamycin or silencing mTOR in ECs of *PGRP-SA* deficient flies brought back normal fat levels (comparison between 20-day old *yw* and PGRP-SA-rapamycin or *PGRP-SA; NP1^ts^>mTOR-RNAi* statistically indistinguishable). This was dependent on 4E-BP however, since fat levels of PGRP-SA; 4E-BP double mutants or *PGRP-SA; NP1^ts^>4E-BP-RNAi* were low and insensitive to rapamycin (comparison between 20-day old PGRP-SA-rapamycin and PGRP-SA; 4E-BP-rapamycin or *PGRP-SA; NP1^ts^>4E-BP-RNAi* is p<0.001 but the latter when compared to 20-day old PGRP-SA or 20-day old 4E-BP gives a n on-significant comparison). This indicated that m TOR/4E-BP signalling in the gut influenced systemic fat levels. Error bars represent standard deviation. Bars represent mean values from three independent experiments.

This suggested that loss of Toll signalling released TOR activity, which dampened 4E-BP’s on lipid metabolism. When TOR activity was inhibited (pharmacologically or through RNAi) in *PGRP-SA* mutants, 4E-BP was able to put a brake on fat wasting and microbiota depletion. Indeed, treatment of PGRP-SA; 4E-BP double mutants or *PGRP-SA; NP1^ts^*>*4E-BP^RNAi^* flies with rapamycin was ineffective in restoring fat levels (Fig. 9) and intestinal CFUs (Fig. S9) indicating that 4E-BP was responsible for regulating fat levels and intestinal microbiota in ECs. As differences in food intake can modulate both lipid reserves as well as lifespan, we measured food intake during a week of observation (15 to 22-day old mated female flies) using a <u>c</u>apillary <u>f</u>eeding assay (CAFE assay, Ja *et al*, 2007). Food intake was statistically indistinguishable between PGRP-SA and 4E-BP single mutants or PGRP-SA; 4E-BP double mutants compared to *yw* controls (Fig. S10).

Long-term intestinal regeneration is important for preserving organ function. In mice, the presence of TLR4 has been linked to increased proliferation of ISCs (Neal et al, 2012). However, the effect of the absence of TLRs has been less clear. Our results show that following infection as well as in the absence of immune challenge loss of Toll signalling blocked long-term epithelial renewal and reduced gut microbiota. Conversely, constitutive Toll in progenitor cells increased EB numbers by increasing ISC proliferation and blocking EBs to differentiate. More work is needed to pinpoint how this mechanism operates.

Infection or constitutive Toll in progenitor cells induced 4E-BP transcription in ECs. 4E-BP has been shown to preserve fat storage under stress conditions while loss of its activity resulted in flies “burning” fat faster (Teleman et al, 2005). Toll-mediated induction of 4E-BP in ECs increased both intestinal and systemic fat levels. Conversely, fat reserves were depleted when Toll was similarly expressed but in a 4E-BP null genetic background or when 4E-BP was silenced in ECs.

Age-dependent loss of fat was also observed in the absence of infection in *PGRP-SA^*seml*^* mutants. Reduction of fat levels was reversed with the use of rapamycin, which targets mTORC1 (reviewed in Laplante and Sabatini, 2012) or by blocking mTOR by RNAi in ECs. This result suggested that absence of Toll signalling elevated mTORC1 activity, which in turn dampened 4E-BP (Hay and Sonenberg 2004). In keeping with this hypothesis *PGRP-SA; 4E-BP* or *PGRP-SA; NP1^ts^>4E-BP^RNAi^* flies treated with rapamycin were unable to restore fat levels. This indicated that systemic fat levels were under 4E-BP control in ECs downstream of Toll. The latter antagonised TOR in the regulation of fat metabolism through 4E-BP.

It is tempting to speculate that in the absence of infection, PGRP-SA recognises parts of the intestinal microbiota and as such activates the Toll pathway in progenitor cells, which in turn keeps 4E-BP activity in enterocytes thus preserving fat levels that are important for normal microbiota population density. Indeed, we have observed that 4E-BP is important for the maintenance of normal CFUs in the gut and we believe that it is the preservation of fat levels through 4E-BP that mediates the maintenance of a normal microbiota population density. In this context, PGRP-SA function resembles intestinal TLR-2, Myd88-mediated responses in the T-cell compartment of the mouse gut (Kubinak et al, 2015). More work is needed to pinpoint, which constituent(s) of the microbiota PGRP-SA may recognise and how the signal is transmitted from the progenitor cells to ECs.

Our data support a model where systemic immunity to infection is directly linked to epithelial renewal and intestinal regulation of fat levels through the evolutionary conserved Toll receptor. This ties together the immune and metabolic aspects of intestinal physiology with long-term epithelial and microbiota homeostasis.

## Materials and Methods

### Fly strains

The genetic backgrounds control strains used in the experiments are yw. We incorporated the *PGRP-SA^*seml*^* mutation (Michel *et al*, 2001) in *esg*^*ts*^>*GAL4* (Buchon et al, 2009). We used the *spz*^*rm*^*7*^^ mutant (de Gregorio *et al,* 2002) and incorporated UAS-CG7923^RNAi^, UAS-9080RNAi and UAS-Toll^RNAi^ in *esg*^*ts*^>*GAL4* and *Su[H]^ts^>GAL4* (Zeng et al, 2010). All RNAi strains were obtained from the Vienna Stock Centre (Dietzl et al, 2007). Thor-lacZ (y’,w*; p{lacW}Thor^K13517^) and Thor^2^ was obtained from Bloomington Stock Centre IN USA (# BL 9558 and # BL 9559 respectively). For Toll constitutive expression we used UAS-Toll10B (Shia *et al,* 2009).

### Infection

To infect flies, Candida albicans (C. albicans) strain was cultured in Sabouraud’s glucose broth (SGB; Oxoid) for 18 hours; cells were harvested by centrifugation (3200 rpm for 5 minutes) and washed in sterile phosphate buffered saline (PBS). Washed fungal cells were again centrifuged and re-suspended in PBS to an optical density of approximately 0.95-1.05 (Thermo Scientific NanoDrop 1000 spectrophotometer). The inoculant containing C. albicans strain was further diluted four-fold in PBS. Similarly, Staphylococcus aureus (S. aureus) NCTC8325-4 was cultured in TSB for 16 hours; cells were harvested by centrifugation (4000 rpm for 7 minutes) and washed in PBS. Cells were then centrifuged and re-suspended in PBS to an optical density of approximately 0.360 and further diluted 1000-fold in PBS for injection. Anaesthetized female flies were infected with 13.2nl of the C. albicans or S. aureus suspensions (or with PBS control), directly injected into the haemolymph through the dorsolateral region of the thorax, using a micro-injector (Drummond Scientific Nanoinject II). The number of viable yeast cells injected per fly was approximately 600, as calculated from plating homogenates of six injected flies, previously ground in SGB medium. Flies were kept at 30°C post-infection for 36 hours and then dissected.

### Gut dissection and immunostaining

For gut imaging, guts from anesthetized flies were dissected in Schneider’s medium and fixed for 30 min in 4% paraformaldehyde (in PBS), rinsed in PBS and then three times washed (5 min each) in wash solution, 0.1% Triton X-100 (Sigma-Aldrich) in PBS. The tissue was blocked for 60 min in blocking solution (0.1% Triton X-100, 2% BSA (Sigma-Aldrich) in PBS and immunostained with primary antibodies overnight at 4°C. Samples were then washed 4 x 5 min at room temperature (RT) In wash solution, incubated with secondary antibodies at RT for 2 hours, washed again as before and were them stained with DAPI 1:1000 (Sigma-Aldrich). Washed guts were mounted in slides with vectorshield mounting media (Vector Laboratories). The following primary antibodies were used: mouse anti-β-galactosidase (40-1a-S, Developmental Studies Hybridoma Bank, Iowa, USA) - 1:1000; goat anti-HRP (123-165-021, Jackson ImmunoResearch Labs. Inc.)-1:500. we used donkey anti-mouse Alexa 568 antibody (Invitrogen) - 1:250 and donkey anti-goat Alexa 568 antibody (Invitrogen) - 1:250.

### Imaging data analysis

Guts were imaged at 20x magnification, and all GFP marked cells (esg > Gal4) co-localised with DAPI small nuclei were counted in an area of approximate size that extended anteriorly from Boundary 2-3 (Buchon N et al; Cell Reports, 2013); plotted values are the number of GFP marked cells per unit area or per total number of DAPI stained cells. Images were analysed using ImageJ software.

### Microbiota analysis

At the indicated age-points six female flies from each of three different vials, each containing approximately 20 flies, were assayed for total microbiota load. The microbiota load was determined both by plating fly extracts and by PCR amplification of the 16S ribosomal RNA gene from DNA obtained from dissected guts. The remaining flies were transferred to fresh vials every two days. Flies were first washed with cold ethanol (70%) and then rinsed in PBS. Flies were homogenized in M.R.S broth, the extract dilutions were then spread on 229 M.R.S agar and the plates were incubated at 30°C. After 48 hours, colonies were counted. For PCR assay, total DNA was extracted from dissected midguts (crop and hindgut were removed) using a G hypodermic needle attached to a homogenizer and using the Cells and Tissue DNA Isolation Kit (NORGEN). The 16S region PCR amplification was carried out for 10ng of each DNA sample, using 50μl reaction mixtures. The primer sequences used were AGAGTTTGATCCTGGCTCAG (16S_27_Fw) and GGTTACCTTGTTACGACTT (16S_1492_Rv). A 482bp fragment from actin was also amplified in these reactions as an internal control. The primer sequences used were CTGGACTTCGAGCAGGAGAT (Act5C3_Fw) and GGTGGCTTGGATGCTTAGAA (Act5C2_Rv). Each reaction mixture contained 0.5μM of each primer, a 200μM concentration of each deoxynucleoside triphosphate (dNTP), and 1μM Phusion Taq polymerase (New England Biolabs). The PCR conditions 236 involved an initial denaturation step at 98°C for 30 secs followed by 35 cycles of 95°C for 30 secs, 65°C for 30 sec, and 72°C for 1 min and ended with an extension step at 72°C for 5 min in a thermocycler (T100 thermal cycler, Bio-Rad). PCR samples were run in a 0.8% agarose gel. The gel was stained with ethidium bromide, visualized and digitally photographed by Alphalimager HP Gel (Alpha 240 Innotech) imaging system. The bands intensities were analysed in Image J.

### Triglyceride measurement

This was done as in Teleman et al, 2005. Briefly, newly hatched L1 larvae were seeded in vials as at density of 50/vial and grown at 25°C to synchronise the culture. Adults at the appropriate age were processed in batches of eight for males and six for females. Only male data are presented (of note that females did not deviate from the results obtained). Samples were processed immediately in homogenisation buffer [0.05% Tween-20 and 2x protease inhibitor (Roche) in H_2_O]. After centrifugation (5000 rpm, 1min) the supernatant was transferred to a new tube and span again (14000 rpm, 3mins at 4°C). To measure triglycerides 80μl of the above supernatant were mixed with 1ml of the Triglyceride Reagent (Thermo-Fisher) and incubated for 10mins at 37°C. Measurements were taken at OD _520_ and compared with a standardization curve. To measure protein levels, 100μl of the final supernatant was combined with 700μl of H_2_O and 200μl of Bio-Rad Protein Assay Reagent and incubated at room temperature for 3mins. Measurements at OD_595_ were compared with a standardization curve.

### Rapamycin feeding protocol

Fly food was microwaved and Rapamycin antibiotic- or ethanol as vehicle control was added to a final concentration of 200 μM (rapamycin is soluble in EtOH, rapamycin was Sigma 37094-10MG). The mix was then added to vials in batches of 10mls. We cultured female flies of the appropriate age (20 flies per v ial) i n vials containing rapamycin o r ethanol treated food. Food was changed every two days for two weeks. When flies reached 20 days of age we sacrificed them and performed downstream experiments (fat level measurements or CFUs).

### <u>Ca</u>pillary <u>Fe</u>eding Assay (CAFE assay)

Food intake was analysed as previously described (Ja *et al*, 2007) with some modifications. 50 flies per genotype were tested. Batches of 10 flies were placed in vials with wet tissue paper at the bottom and a capillary food source containing a blue dye. Feeding was monitored for 8 hours (light ON) and 1 hour (light OFF). Feeding amount was recorded every 1 hour and the capillaries were replaced every 2 days.

### Statistical analysis

Data was analysed using GraphPad Prism 6 or R. First a D’ Agostino and Pearson omnibus Normality test was conducted. If the data was found to fit a normal distribution, parametric tests were used, first ANNOVA and then a Tukey’s multiple comparisons test. For the Thor-LacZ count data in the cases that did not fit the normal distribution we conduct Kruskal-Wallis test for the followed by the Dunn’s multiple comparisons test to clarify the significance. R was used to analyse the GFP count data, it was fitted to a generalised linear model using a quasi-Poisson regression and then ANNOVA and Tukey’s multiple comparisons tests were employed to look for significance. For qPCR gene expression data was standardized by series of sequential corrections, including log transformation, mean centring, and autoscaling (Willems et al, 2008).

## Acknowledgements

We would like to thank the Vienna and Bloomington Stock Centres, as well as Bruno Lemaitre for fly stocks and the Iowa Hybridoma Bank for antibodies. This work was funded by ERC Consolidator Grant 310912 and BBSRC Responsive Mode Grant BB/P005691/1 (both to PL).

---

**Figure S1. Toll signalling endorses ISC but not EB cell proliferation. (A)** Constitutive Toll signalling did not induce EB proliferation. *Su(H)^ts^>Toll10B* flies, which express the constitutively active Toll receptor variant T oll10B (green channel) did not activate proliferation as s hown by the lack o f staining for pH3 (red channel). All cells were stained with DAPI (blue channel). **(B)** Silencing Toll in ISCs with a *esg*^*ts*^>*UAS-Toll-RNAi* configuration prevented ISC proliferation 36h following *C. albicans* infection (lower panel) compared to control flies (upper panel).

**Figure S2. Silencing expression of Toll in EBs reduces their number. (A)** RNAi knockdown of the Toll receptor specifically in EBs [Su(H)-Gal4] of the adult gut leads to reduction in cell number and alters their morphology, where the cells generally remain rounded, rarely adopting the characteristics elongated and irregular shape often observed with ISCs/EBs (compared cells marked with white asterisks). CG7923 RNAi line was randomly chosen from the VDRC library and used as an internal control for the RNAi effect. EBs marked with GFP (green) and all nucleic stained with DAPI (grey) in 20-day old flies. **(B)** Quantification showed significant EB loss in the absence of Toll (95% confidence interval is displayed).

**Figure S3. Proliferation of ISCs is absent from the intestinal epithelium of *PGRP-SA^*seml*^* mutants.**

**(A)** Guts from 20-day old *ywPGRP-SA^*seml*^; esg^ts^>GFP* females were stained with anti-GFP (green channel), DAPI (blue channel) and anti-pH3 (red channel). This is a representative sample from n=10.

**(B)** In contrast, *yw* flies were found to have a significantly higher number of ISCs proliferating (compare red channels).

**Figure S4. Loss of *spz* results in the reduction of ECs.** Staining of *esg*^*ts*^>*GFP; spz*^*rm*^*7*^^ or *esg*^*ts*^>*GFP; spz*^*rm*^*7*^^/+ with anti-GFP (to mark progenitor cells, upper panels) and DAPI (to distinguish ECs, lower panels) in 20-day intestines. The homozygous *spz*^*rm*^*7*^^ flies exhibited significantly lower numbers of ECs than heterozygous *spz*^*rm*^*7*^^ flies (p<0.05, unpaired t-test). Each dot in the graph represents the average number of ECs from 15 guts as measured placing a square always the same size at a random point on the surface plane of the anterior midgut epithelium.

**Figure S5. Constitutive expression of Toll in progenitor cells rescues ISC numbers in the absence of PGRP-SA. (A)** Expression of Toll10B in progenitor cells (*esg*^*ts*^>*Toll10B*) in a genetic background mutant for *PGRP-SA* recovers ISCs cell numbers; ISCs are labelled in red (HRP staining) while ISCs and EBs are labelled in green (GFP) in 20-day old flies. **(B)** This can be verified comparing the *yw* genetic background with the *PGRP-SA^*seml*^*; *esg*^*ts*^>*Toll10B* flies where ISC numbers were statistically indistinguishable; (** p<0.01).

**Figure S6. Loss of Toll signalling results in loss of intestinal microbiota. (A)** Semi-quantitative PCR of 16S rRNA shows an age-dependent reduction in total intestinal microbiota when comparing *yw* to *PGRP-SA^*seml*^* or *Dif*^*1*^ but no reduction in the internal control (actin). **(B)** Quantification showed a significant reduction in the quantities of 16S.

**Figure S7. Loss of PGRP-SA function reduces lifespan.** Both female and male flies with a deficiency in PGRP-SA function (carrying the *seml* mutation) have a significantly reduced lifespan. Median lifespan of control *yw* females was 58 days vs. 20 days for *seml* females while median lifespan of control *yw* males was 41 days vs. 18.5 for *seml* males.

**Figure S8. Toll signalling regulates intestinal triglyceride levels through the TOR/4E-BP axis.**

Following *C. albicans* infection, intestinal fat levels increase (comparison between yw and yw-*C. albicans* p<0.001). The presence of d4E-BP ensures that intestinal fat levels are not “burnt” as fast during infection since infected Thor mutants (4E-BP-*C. albicans*) or f lies with silenced 4E-BP in the ECs (*NP1^ts^>4E-BP-RNAi*) have significantly less fat (comparison between yw-*C. albicans* and 4E-BP or NP1ts>4E-BP-RNAi p<0.001). Toll overexpression in progenitor cells through *esg-GAL4* increases intestinal fat levels (comparison between *yw-C. albicans* and Toll-10B is non-significant but between yw and Toll-10B p<0.001). However, in a genetic background mutant for 4E-BP, Toll overexpression does not have an effect (comparison between *4E-BP-C. albicans* and Toll non-significant). Error bars represent standard deviation. Bars represent mean values from three independent experiments.

**Figure S9. Toll signalling regulates intestinal microbiota levels through the TOR/4E-BP axis.** 20-day old *yw* flies display significantly more intestinal CFUs than *PGRP-SA^*seml*^* mutants (p<0.01). Loss of CFUs can be rectified by treating *PGRP-SA^*seml*^* with rapamycin. However, this is dependent on 4E-BP since treatment with rapamycin of a double *PGRP-SA; 4E-BP* mutant (or flies mutant for *PGRP-SA* and with 4E-BP silenced in ECs) have CFUs at the level of *PGRP-SA^*seml*^*.

**Figure S10. Food intake is not influenced by lack of either PGRP-SA or/and 4E-BP function.** Food consumption was measured by the CAFE method in mated females (as they had a better life expectancy in PGRP-SA mutants see Fig. S6) from day 15 to day 22 of adulthood; n=5 vials (of 10 flies each) per genotype; no comparison was statistically significant (p>0.05, unpaired t-test).

## Notes

#### Summary of Updates

The lipid metabolism experiments (Figs 8 and 9 and Supplemental Figs S8, S9 and S10) have been expanded; the manuscript includes lifespan data from PGRP-SA mutants (Fig S7).

